# Activation of Store-Operated Calcium Entry and Mitochondrial Respiration by Enterovirus 71 Is Essential for Efficient Virus Replication

**DOI:** 10.1101/2023.12.20.572621

**Authors:** Bang-Yan Hsu, Ya-Hui Tsai, Ta-Chou Weng, Szu-Hao Kung

## Abstract

Enterovirus (EV) infections disrupt cellular calcium (Ca^2+^) homeostasis. The EV protein 2B is localized to the endoplasmic reticulum (ER) and causes depletion of ER Ca^2+^ stores. This depletion coincides with a substantial increase in cytosolic Ca^2+^ levels driven by extracellular Ca^2+^ influx. However, the precise mechanism underlying this influx remains elusive. In the present study, we demonstrated that EV71 infections induce store-operated Ca^2+^ entry (SOCE) by activating the Ca^2+^ sensor stromal interaction molecule 1 (STIM1), which subsequently interacts with Orai1, a plasma membrane (PM) Ca^2+^ channel. This finding was supported by confocal imaging, which revealed that STIM1, typically localized in the ER, becomes active and colocalizes with Orai1 at the PM in EV71-infected cells. Pharmacological inhibition of the STIM1–Orai1 interaction and knockdown of either STIM1 or Orai1 significantly reduced virus-induced cytosolic Ca^2+^ levels and viral replication. Global transcriptome analysis revealed that differentially expressed genes are primarily associated with the mitochondrial electron transport chain (ETC) upon SOCE activation, contributing to enhanced ATP generation and oxygen consumption. This increase in mitochondrial Ca^2+^ levels is correlated with the mid-stage of virus infection. Furthermore, we demonstrated that high levels of mitochondrial Ca^2+^ influx led to apoptotic cell death favoring viral release at the late stage of virus infection. Finally, SOCE-dependent EV replication was observed in a mouse intestinal organoid culture, a more physiologically relevant cell system. Our results provide valuable insights into the mechanism through which EV infections induce SOCE-mediated spatial and temporal control of Ca^2+^ signaling, substantially affecting the virus life cycle.

**IMPORTANCE:** Host cell Ca^2+^ signals play crucial roles in various steps of virus life cycles, including entry, replication, and exit. EV requires increased cytosolic Ca^2+^ levels for efficient replication, but the precise mechanisms underlying the association between Ca^2+^ levels and EV replication remain elusive. Using EV71 as a model virus, we demonstrated that EV71 infection elevated cytosolic Ca^2+^ levels through store-operated Ca^2+^ entry activation and progressive Ca^2+^ mobilization to mitochondria. This led to the upregulation of electron transport chain activity, which is essential for efficient virus replication and apoptotic cell death, facilitating viral release during the mid and late stages of the infectious cycle, respectively. These findings substantially enhance the understanding of how EVs co-opt host cell mechanisms to promote their own life cycle. STIM1 and Orai1 may be novel targets for broad-spectrum host-directed therapeutics against EVs and other viruses that employ similar replication mechanisms.

## INTRODUCTION

Enteroviruses (EVs) are nonenveloped, positive-strand RNA viruses belonging to the genus *Enterovirus* of the family *Picornaviridae*. EVs consist of many human pathogens, including poliovirus, coxsackie A and B viruses, echoviruses, numbered EVs, and rhinoviruses [1]. Diverse EVs can cause a wide array of clinical manifestations ranging from mild illnesses to more severe and even life-threatening diseases, such as myocarditis, meningitis, encephalitis and acute paralysis. Among EVs, EV71 (or EV-A71) infections have caused several outbreaks of hand, foot and mouth disease, with a notable proportion of infected infants and young children manifesting severe neurological complications and even consequent death, mainly in the Asia-Pacific region [1, 2]. However, currently, there are no approved antiviral agents available against EV infections.

Intracellular calcium (Ca^2+^) concentration plays a crucial role in regulating various cellular functions involved in physiological and pathological states. Viruses, as obligate intracellular parasites, have developed strategies to manipulate Ca^2+^ signaling and Ca^2+^-dependent processes to promote their own life cycles [3, 4]. During the course of EV infection, a gradual increase in the cytosolic Ca^2+^ level was observed, due partly to the increased permeability of the endoplasmic reticulum (ER) and the consequent depletion of Ca^2+^ stores in the ER, causing outflow of stored Ca^2+^ into cytoplasm [5, 6, 7, 8, 9]. In addition, the influx of extracellular Ca^2+^ into EV-infected cells is thought to contribute to increased cytosolic Ca^2+^ concentrations as well [7, 10]. Disruption in Ca^2+^ homeostasis caused by EV infection is pivotal in viral replication. The use of Ca^2+^ chelators was reported to substantially reduce the production of viral proteins and progeny viruses [11].

Studies have demonstrated that EV 2B proteins play a pivotal role in increasing cytosolic Ca^2+^ levels. These proteins act as viroporins, a type of virus-encoded pore-forming protein [12, 13, 14]. EV 2B proteins are small, integral membrane polypeptides consisting of approximately 99 amino acids. They contain two hydrophobic regions connected by a short stem-loop. One of these hydrophobic regions is likely to form an amphipathic alpha helix, whereas the other is likely to form a complete hydrophobic helix. This helix–loop–helix motif of 2B is believed to be responsible for its ability to form transmembrane pores through oligomerization [12]. In EV-infected cells, 2B proteins localize in membranes derived from the ER and Golgi apparatus. The pore-forming activity of 2B in these organelle membranes results in a decrease in Ca^2+^ stores in the ER and Golgi apparatus [10, 12], and is thus functionally associated with viral replication and release [8, 10, 15]. The single expression of EV 2B is sufficient to recapitulate the changes in Ca^2+^ homeostasis observed in EV-infected cells [7, 8]. However, the molecular mechanism underlying the EV 2B–induced increase in cytosolic Ca^2+^ levels that supports viral replication and release remains elusive.

The coordinated regulation of Ca^2+^ release from the ER and subsequent entry of Ca^2+^ across the plasma membrane (PM) to replenish ER stores is referred to as store-operated calcium entry (SOCE). SOCE is a homeostatic cellular mechanism that helps maintain appropriate Ca^2+^ levels in the ER to facilitate Ca^2+^-mediated signaling [16]. Two protein families, stromal interaction molecules (STIMs) and Ca^2+^ release-activated Ca^2+^ modulators (Orais), are responsible for activating SOCE [16, 17]. STIM proteins act as sensors for Ca^2+^ levels in the ER. When Ca^2+^ is depleted in the ER [18], STIM proteins aggregate into multiple puncta that translocate close to the PM. Orai, an essential pore-forming component of SOCE, translocates to the same STIM-containing structures during Ca^2+^ depletion in the ER and mediates extracellular Ca^2+^ entry. To date, three isoforms of Orai (1–3) and two isoforms of STIM (1 and 2) have been identified in mammals. STIM1 levels are higher than STIM2 levels in most tissues. Moreover, STIM1 and Orai1 are sufficient for the generation of Ca^2+^-release-activated Ca^2+^ currents, making Orai1 the primary channel activated by Ca^2+^ store release and STIM1 activation [19].

Given that EV 2B protein resides in the ER and that EV-infected or 2B-expressing cells experience an influx of extracellular Ca^2+^, we investigated whether the ER–PM connection is involved in the increased cytosolic Ca^2+^ levels in the presence of EV 2B, i.e., SOCE activation, is involved in increased cytosolic Ca^2+^. In the present study, we used EV71, known to cause disrupted Ca^2+^ homeostasis, as a model [20, 21]. We demonstrated that EV-infected cells activate SOCE, with this activation mediated by the interaction between STIM1 and Orai1. To understand the downstream Ca^2+^ signaling events that support viral replication, we performed bulk RNA-sequencing (RNA-seq) to analyze the transcriptome of EV71-infected cells and the transcriptional reprogramming of a specific SOCE inhibitor, AnCoA4. We determined that SOCE activation led to an increase in mitochondrial Ca^2+^ levels, thereby promoting ATP production and cellular respiration by transcriptionally upregulating a panel of genes encoding mitochondrial electron transport chain (ETC) complexes at 6 h postinfection (p.i.). Treatment with oligomycin, an inhibitor of ATP production, reduced viral replication in a dose-dependent manner, indicating that increased ATP production is required for efficient EV71 replication. At 12 h p.i., Ca^2+^ influx led to mitochondrial Ca^2+^ overload, resulting in the activation of intrinsic apoptosis that facilitated virus egress and pathogenesis. This SOCE and mitochondria Ca^2+^-dependent viral replication phenomenon was also noted in other EV serotypes. Finally, we observed SOCE-dependent upregulation of ETC complexes and EV replication in a mouse intestinal organoid culture, a physiologically-relevant model known for its prior uses in studies of cell Ca^2+^ flux and mitochondrial respiration [22, 23, 24].

## MATERIALS AND METHODS

### Cells and Viruses

Rhabdomyosarcoma (RD) cells (ATCC, CCL-13) and HeLa cells (ATCC, CCL-2) were cultured as described elsewhere [25]. Human EV stocks used in this study included EV71 (strain BrCr), coxsackievirus A16 (CVA16), CVB3 and echovirus serotype 30 (Echo30) as reported previously [25]. A mouse-adapted EV71 MP4 strain [26] was provided by Dr. Jen-Ren Wang (National Cheng Kung University, Taiwan). All virus stocks were titrated on RD cells by 50% tissue culture infectious dose (TCID_50_) assay.

### Ca^2+^ fluorescence dyes assay

RD and HeLa cells were infected with EV71 at a specified multiplicity of infection (MOI) in 24-well plates for the indicated durations. Cells were incubated with the cytosolic Ca^2+^ dye Fluo-8 at 1 µM or the mitochondrial Ca^2+^ dye Rhod-2 AM at 3 µM, both for 30 min at 37°C. The emitted fluorescence resulting from Fluo-8 or Rhod-2 AM staining was observed using a fluorescence microscope (Leica DM6000B) with a 488-nm or 543-nm filter, respectively. The fluorescence intensity was quantified using MetaMorph software (Molecular Devices, Downington, PA).

### Ectopic expressions of STIM1 and Orai1 for confocal analysis

Plasmid encoding enhanced green fluorescent protein (EGFP)-labeled STIM1 (pEGFP-STIM1) and monomeric red fluorescent protein (mOrange)-labeled Orai1 (pmOrange-Orai1) were provided by Dr. Meng-Ru Shen (National Cheng Kung University, Taiwan). The colocalization of STIM1 and Orai1 was observed by confocal microscopy. See Text S1 for details.

### RNA-seq

RD cells were infected with EV71 at an MOI of 0.5 for 6 h in the presence of DMSO or 20µM AnCoA4. RNA extraction was performed using RNeasy Kits (Qiagen, Hamburg, Germany), and RNA-seq libraries were prepared using the Universal Plus mRNA Seq Preparation kit (Tecan). RNA-seq was performed using the NovaSeq 6000 platform (Illumina, San Diego, CA) in accordance with the manufacturer’s protocol, with 2 × 150-bp paired-end sequencing. Reads were aligned to the GRCm38 build of the human genome by using CLC Genomics Workbench v10 (Qiagen).

### Differential expression analysis

The expression level of each gene was examined by fragment length, determined by read counts, and normalized using the fragments per kilobase of transcript per million mapped fragments variation method. Genes with significant changes in mRNA levels were identified on the basis of a false discovery rate cutoff of 0.05 and a fold change of 2. Gene set enrichment analysis was conducted using the Database for Annotation, Visualization, and Integrated Discovery (DAVID, https://david.ncifcrf.gov/) [27]. Biological processes with corrected *p*-values of *≤*0.05 were considered significantly enriched pathways.\

### Measurement of cellular respiration

Cellular respiration was measured using the Seahorse XF Cell Mito Stress Test Kit (Agilent, Santa Clara, CA) in accordance with the manufacturer’s instructions. HeLa cells seeded in XFe 24-well microplates were infected with EV71 stock at an MOI of 5 for 6 h. The microplates were placed in a non-CO_2_ incubator at 37°C for 1 h prior to measurement. Cellular respiration was analyzed using sequential injections of 2 µM oligomycin (an ATP synthase blocker), 1 µM carbonyl cyanide-p-trifluoromethoxyphenylhydrazone (FCCP, an activator of the uncoupling of the inner mitochondrial membrane), and 0.5 µM rotenone/antimycin A (inhibitors of complexes I and III). The oxygen consumption rate (OCR) was determined by Seahorse XFe24 Extracellular Flux Analyzer (Agilent).

### Cultivation, infection and imaging of crypt organoids

Mouse intestinal stem cells were isolated following a method documented [28] and it is reported in the Text S1. The preparation and culture of mouse intestinal organoid followed a protocol described previously with certain modifications [22]. Briefly, mouse crypt organoids were seeded in differentiation medium consisting of Advanced DMEM/F12 (Invitrogen) containing growth factors as previously reported [28]. Crypt organoids were mixed with Matrigel (BD Bioscience) and plated in 12-well plates. The medium was changed every 2 days for 10-14 days. Differentiated mouse intestinal organoids were sheared mechanically by pipetting to enable sufficient apical exposure to virus inoculums. The organoids were inoculated with 200 µL EV71 MP4 strain (4 *×* 10^8^ TCID_50_/mL) for 2 h at 37°C. After the incubation, the organoids were washed with PBS and organoids were embedded in Matrigel with the differentiation medium at 37 °C for 48 h. The images were acquired by confocal microscopy (LSM880, Zeiss). See Text S1 for details.

## RESULT

### EV71 infection induced Ca^2+^ influx that is required for efficient virus replication

Previous studies have demonstrated that infection with poliovirus or coxsackievirus B increases cytosolic Ca^2+^ levels [5, 6]. In the current study, we used EV71 as a model virus due to its known activities in triggering Ca^2+^ flux and Ca^2+^-dependent signaling [20]. To investigate the Ca^2+^ flux and its impact on EV71 replication in a single-cycle infection, RD cells were infected with EV71 at a lower or a higher dose, i.e., an MOI of 0.5 or 5, respectively, for 12 h. Our results revealed that the cytosolic Ca^2+^ levels significantly increased starting at 4 or 6 h p.i. upon 0.5- or 5-MOI infection, respectively, and continued to rise throughout the infection, as indicated by staining with the fluorescent Ca^2+^-binding dye Fluo-8 (Fig. 1A, 1C). To further identify the source of Ca^2+^, we added intracellular and extracellular Ca^2+^ chelating agents, BAPTA-AM (referred to as BAPTA hereafter) and EGTA, respectively, both at non-cytotoxic concentrations (Fig. S1). At 0.5 MOI infection, both treatments resulted in a marked reduction in cytosolic Ca^2+^ levels between 6 and 12 h p.i. (Fig. 1A, S2), indicating that at least part of the increased cytosolic Ca^2+^ originated from outside the infected cells in this condition. Viral titers were also reduced upon the treatments at all points tested during the infection (Fig. 1B). On the other hand, infection at 5 MOI led to a significant reduction in the cytosolic Ca^2+^ only at 4 and 6 h p.i., and Ca^2+^ raised to the levels indistinguishable from that of the controls for the remaining infection period (Fig. 1C). A similar trend of virus replication kinetics in HeLa cells upon the Ca^2+^ chelator treatments was observed (Fig. S3). In line with the finding, viral replication was significantly delayed at 4 and 6 h p.i. by the treatment of the Ca^2+^ chelating agents but not thereafter. This implies that EV71 infection at 5 MOI resulted in the induction of cytosolic Ca^2+^ to the high levels that are not susceptible to the treatments of the Ca^2+^-chelators late in the infection (9 and 12 h p.i.). Together, these data indicate that extracellular Ca^2+^ influx contributed to the increase in cytosolic Ca^2+^ observed during EV71 infection and that the increase in cytosolic Ca^2+^ level is correlated with the progression of viral replication, consistent with that for other EVs [5, 6].

**FIG 1.**
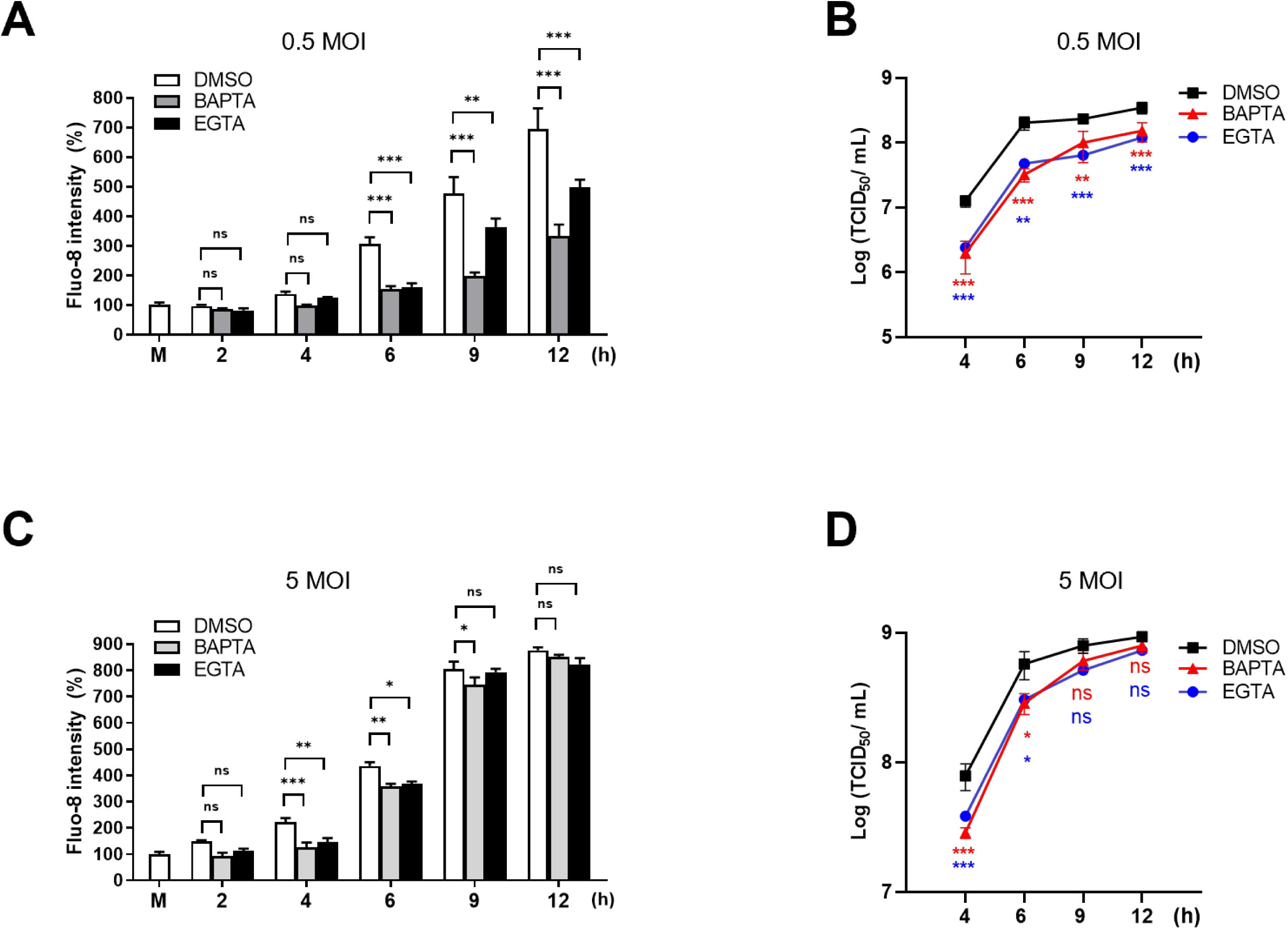
EV71 infection induced elevated cytosolic Ca^2+^ levels that correlated with the progression of viral replication. RD cells were mock-infected (M) or infected with EV71 stock at an MOI of 0.5 (A, B) or 5 (C, D) for the indicated times in the presence of BAPTA (10 µM), EGTA (1.8 mM), or DMSO control. (A, C) Cells were stained with Fluo-8 for 30 mins. Fluorescence intensities were quantified relative to the mock-infected controls. (B, D) Cell lysates and supernatants were collected at the indicated times. Total virus titers were determined by TCID_50_ assay (See Text S1). Data represent mean values from experiments conducted in triplicate, with error bars indicating standard deviations (SDs). \*\*\**p* < 0.001, \*\**p* < 0.01, \**p* < 0.05 versus the DMSO control.

### Orai1 and STIM1 are essential during active replication of EV71

Several Ca^2+^ channels or transporters on the PM mediate Ca^2+^ influx from outside the cells [29]. Among them, only SOCEs are activated by the link between ER-resident STIM1 and PM-resident Orai1. EV 2B localizes in the ER and reduces ER Ca^2+^ levels by forming transmembrane pores [30]. Thus, we hypothesized that SOCE machinery contributes to Ca^2+^ influx extracellularly through the PM, resulting in the observed increase in cytosolic Ca^2+^ levels in the EV71-infected and 2B-expressing cells.

To validate this hypothesis, we first investigated EV71 infection at 6 h p.i., a time point that the increased cytosolic Ca^2+^ and viral replication were most sensitive to the Ca^2+^ chelating agents (Fig. 1B, D), upon the treatment of a specific SOCE blocker AnCoA4 [31]at 10- and 20-µM in RD cells. Dose-dependent reduction in viral titers was found, with viral replication reduced for about a log by 20 µM AnCoA4 (Fig. 2A). For infection with 0.5 MOI, we observed that levels of cytosolic Ca^2+^ were significantly reduced from 6 h to 12 h p.i., similar to those treated by Ca^2+^ chelation (Fig. 1A, 2B). The viral titers were significantly reduced at all time points during the infection period (Fig. 2C) For cells infected with EV71 at 5 MOI, on the other hand, a significant reduction in cytosolic Ca^2+^ and viral titers by AnCoA4 were limited to 4 h and 6 h p.i., and the antiviral effects significantly attenuated with viral titers reaching comparable levels between the treated-and control groups at 9 h and 12 h p.i. (Fig. 2D, E). This finding is consistent with those treated with the Ca^2+^ chelating agents (Fig. 1, S3). A similar trend of virus replication in HeLa cells upon AnCoA4 treatment was noted (Fig. S3) To identify the stage of viral replication promoted by SOCE, we performed a time-of-addition experiment with the treatment of AnCoA4 during distinct stages of virus infection (Fig. 2F, G). Based on the total virus titer determined from each condition, we concluded that AnCoA4 treatment during 3-6 h after viral adsorption, but not other stages tested, displayed significant inhibition on virus replication.

**FIG 2.**
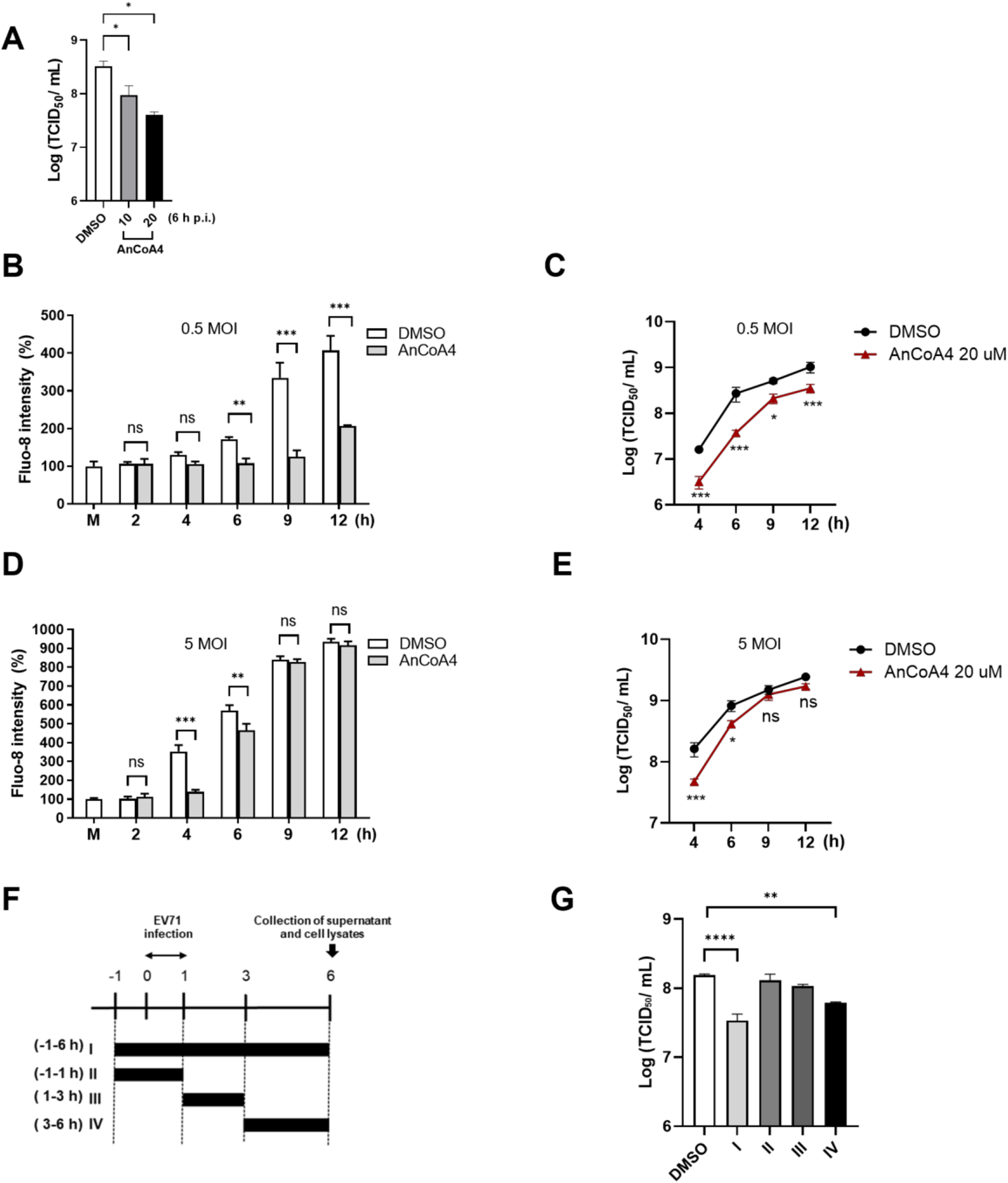
A specific SOCE blocker AnCoA4 reduced virus-induced cytosolic Ca^2+^ levels and viral replication. (A) RD cells were infected with EV71 at an MOI of 0.5 for 6 h in the presence of DMSO control or 10 and 20 µM AnCoA4. RD cells were mock-infected (M) or infected with EV71 stock at an MOI of 0.5 (B, C) or 5 (D, E) for the indicated times in the presence of 20 µM AnCoA4 or DMSO control. (B, D) Cells were stained with Fluo-8 for 30 mins. Fluorescence intensities were quantified relative to the mock-infected controls. (C, E) Cell lysates and supernatants were collected at the indicated time. Total virus titers were determined by TCID_50_ assay. (F) Timing of EV71 inhibition by AnCoA4 with the schematic representation. AnCoA4 was added to RD cells at the indicated times relative to viral infection at an MOI of 0.5, with the initiation (0 h) and completion (1 h) of viral adsorption indicated. The bars indicate the periods with AnCoA4 treatment. Viral supernatants and cell lysates were collected at the end of 6 h for all treatment conditions. (I) Whole: -1 to 6 h; (II) entry: -1 to 1 h; (III) early: 1–3 h; (IV) viral replication: 3-6 h. (G) Total virus titers were determined by TCID_50_ assay. Data represent mean values from experiments conducted in triplicate, with error bars indicating standard deviations (SDs). \*\*\**p* < 0.001, \*\**p* < 0.01, \**p* < 0.05 versus the DMSO control.

To gain more insights into the role of SOCE in EV71 infection, we used lentivirus-mediated RNA interference to generate stable STIM1- and Orai1-knockdown cells, and the efficient knockdown effects were validated (Fig. S4). Knockdown of either STIM1 or Orai1 resulted in marked reductions in EV71-induced cytosolic Ca^2+^ at 6 h p.i. and 12 h p.i., compared with that of the scramble control (Fig. 3A). Likewise, levels of viral protein and viral titers were greatly reduced in the STIM1- and Orai1-knockdown cells over the scramble controls, at both 6 h p.i. and 12 h p.i. (Fig. 3B, C).

**FIG 3.**
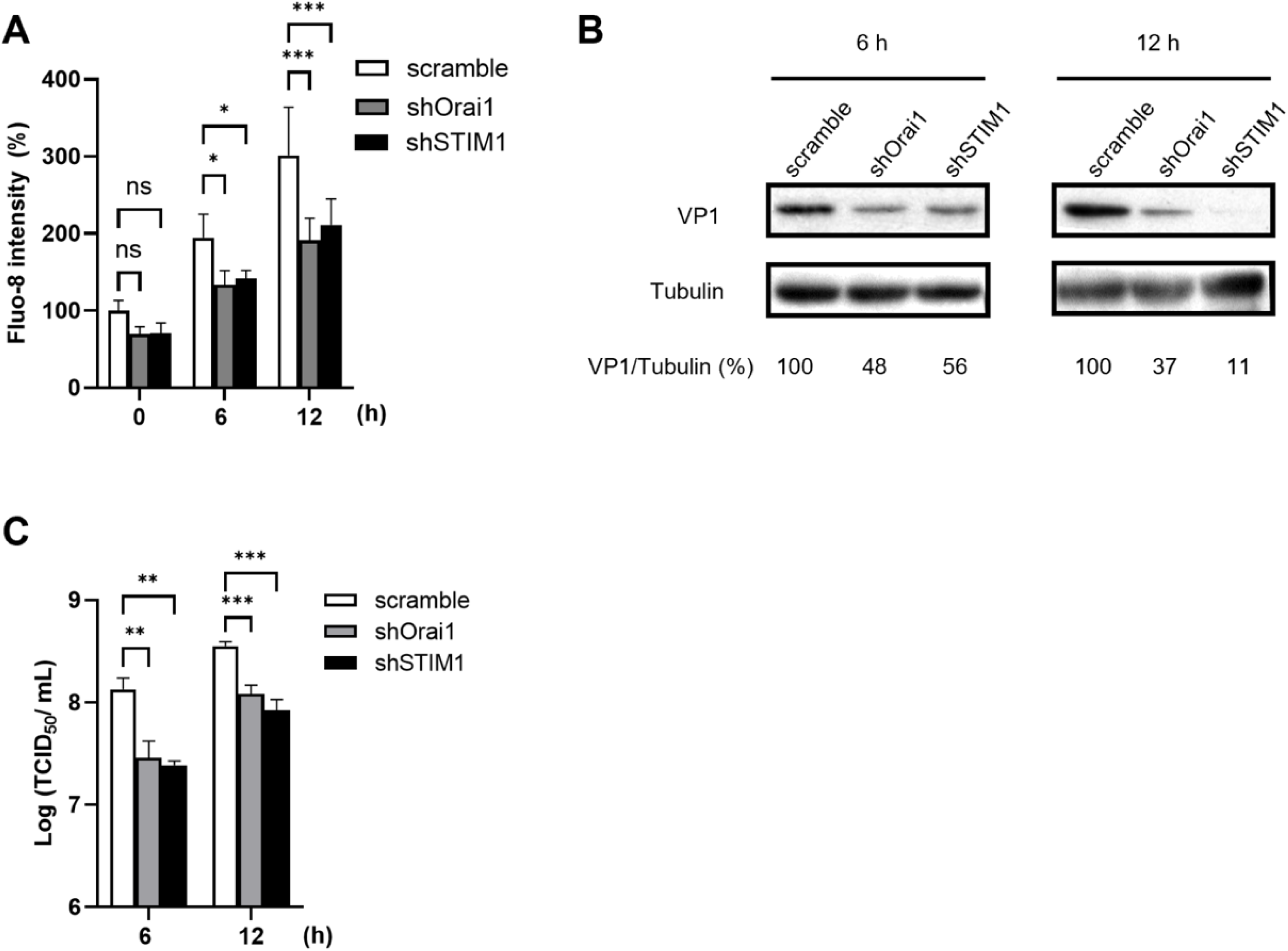
Both Orai1 and STIM1 are essential during active replication of EV71. HeLa cells were transduced with lentivirus–shScramble, –shOrai1 or –shSTIM1 48 h prior to infection with EV71 at 0.5 MOI for 6 h and 12 h. (A) Cells were stained with Fluo-8 for cytosolic Ca^2+^ measurement. (B) Cell lysates were harvested and levels of viral VP1 protein and tubulin (the internal control) were measured by immunoblot analysis. The percentage shown below each lane represents the intensity of viral VP1 relative to that of tubulin. (C) Total virus titers were determined by TCID_50_. The experiments were conducted in triplicate, and shown as means ± SDs. \*\*\**p* < 0.001,\*\**p* < 0.01, \**p* < 0.05 versus the scramble control.

Subsequently, we generated HeLa cells ectopically coexpressing STIM1 and Orai1 labeled with EGFP and mOrange, respectively (Text S1). Transfection of EV71 2B was conducted because single expression of EV 2B expression was sufficient to induce the influx of extracellular Ca^2+^ and to release Ca^2+^ from ER stores [10]. Thapsigargin (TG) was used as a control as it is known to induce ER Ca^2+^ store depletion and activate SOCE by blocking the sarco/ER Ca^2+^-ATPase pump [32]. Whereas the mock-infected and empty-plasmid cells exhibited minimal levels of STIM1-Orai1 colocalization, the infected HeLa-Orai1-STIM1 cells (Fig. 4A) or the cells ectopically expressing viral 2B (Fig. 4B, D) exhibited increased colocalization of STIM1 and Orai1, consistent with that of the TG-treated cells (Fig. 4A, B). Moreover, the colocalization of STIM1 and Orai1 caused by EV71 infection was substantially inhibited by the treatment of AnCoA4 (Fig. 4A). In all cases, colocalization between EGFP-STIM1 and mOrange-Orai1 was quantified by calculation of Mander’s coefficient [33] (Fig 4C, D).

**FIG 4.**
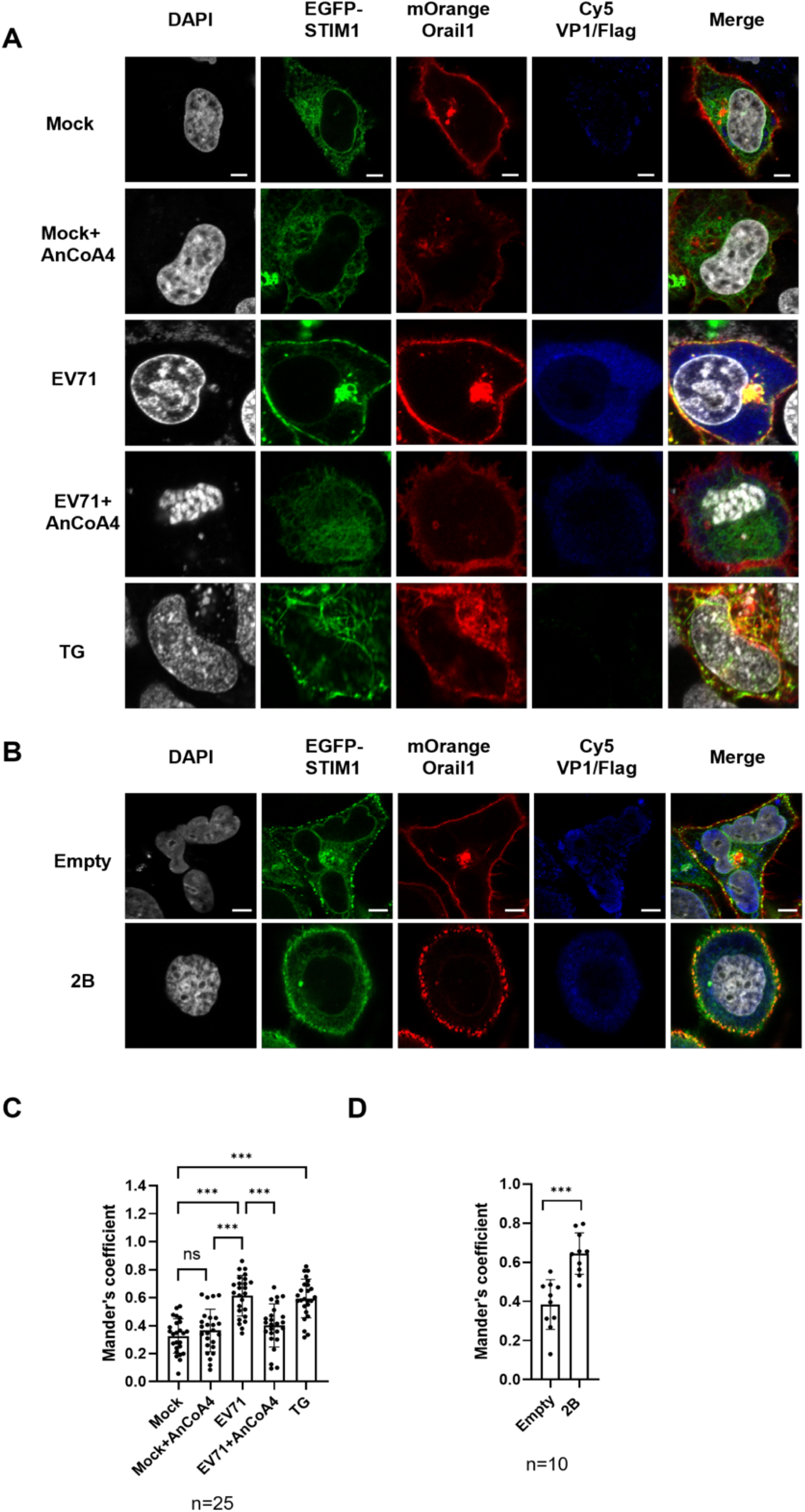
EV71 infection and viral 2B expression induced enhanced colocalization of STIM1 and Orai1. (A) HeLa-STIM1-Orai1 cells coexpressing EGFP–STIM1 (green) and mOrange–Orai1 (red) were mock-infected or infected with EV71 at an MOI of 1 for 9 h in the absence or presence of AnCoA4 at 20 µM. Treatment with 5 µM thapsigargin for 15 mins served as a positive control for SOCE activation. (B) HeLa-STIM1-Orai1 cells were also transfected with pCMV-Flag-2B (2B) plasmid for 48 h. EV71 VP1 (A) or 2B (B) was detected using the anti-EV71 primary Ab or anti-Flag Ab, respectively, followed by the Cy5-labeled secondary antibody (blue) . Colocalization of EGFP–STIM1 and mOrange-Orai1 is presented in yellow. Cell nuclei were stained with DAPI (white). (C, D) The colocalization between EGFP–STIM1 and mOrange-Orai1 was measured using Mander’s coefficients. Each bar represents the mean Mander’s coefficient for a specific experimental condition, reflecting the degree of colocalization between Orai1 and STIM1. Images were acquired using a Zeiss LSM880 confocal microscope. The scale bar represents 5 µM. \*\*\**p* < 0.001,\*\**p* < 0.01, \**p* < 0.05 versus the mock-infected or EV71-infected group.

Taken together, these data indicated that EV71 infection activated the STIM1-Orai1 machinery, resulting in increased cytosolic Ca^2+^, which is essential for active virus replication but not during the entry or initiation of replication.

### Transcriptome profiling identified mitochondrial ETC genes differentially expressed upon the SOCE activation

The well-known SOCE-NFAT axis in Ca^2+^ signaling [31, 34] was found to be independent of EV71 replication (Fig. S5). To investigate the global transcript alterations upon EV71 infection alone and infection together with AnCoA4 treatment, we performed RNA-seq in RD cells at 6 h p.i., a time point that AnCoA4 exerted strong antiviral activity (Fig. 2C, E) and compared the differentially expressed genes. A total of 128 genes with a fold change of >2 were identified (Fig. S6). Gene set enrichment analysis revealed that enriched functional clusters were significantly altered in both groups. These were primarily associated with the mitochondrial ETC (Table 1, Fig. 5A), which consists of four complexes (CI through CIV) that couple reduction–oxidation reactions, creating an electrochemical gradient that leads to the generation of ATP. The identified genes encode mitochondrial NADH dehydrogenases (MT-NDs), cytochrome B (CYB), and cytochrome c oxidase subunit I (COI), which belong to mitochondrial ETC CI, CIII, and CIV, respectively (Table 1); in all cases, EV71 infections upregulated their RNA levels for > 2-fold while the infection with AnCoA4 treatment downregulated those for > 2-fold. We next validated the results through RT-qPCR, which revealed a similar trend of alterations in ND4, ND6, CYB, and COI transcripts following EV71 infection in the absence or presence of AnCoA4 treatment (Fig. 5B). It was noted that the mitochondrial ETC RNA levels from mock-infected cells upon DMSO- or AnCoA4 treatment were not altered, suggesting that cells with SOCE activation are sensitive to AnCoA4 inhibition while those with the basal SOCE levels are not (Fig. 5B).

**FIG 5.**
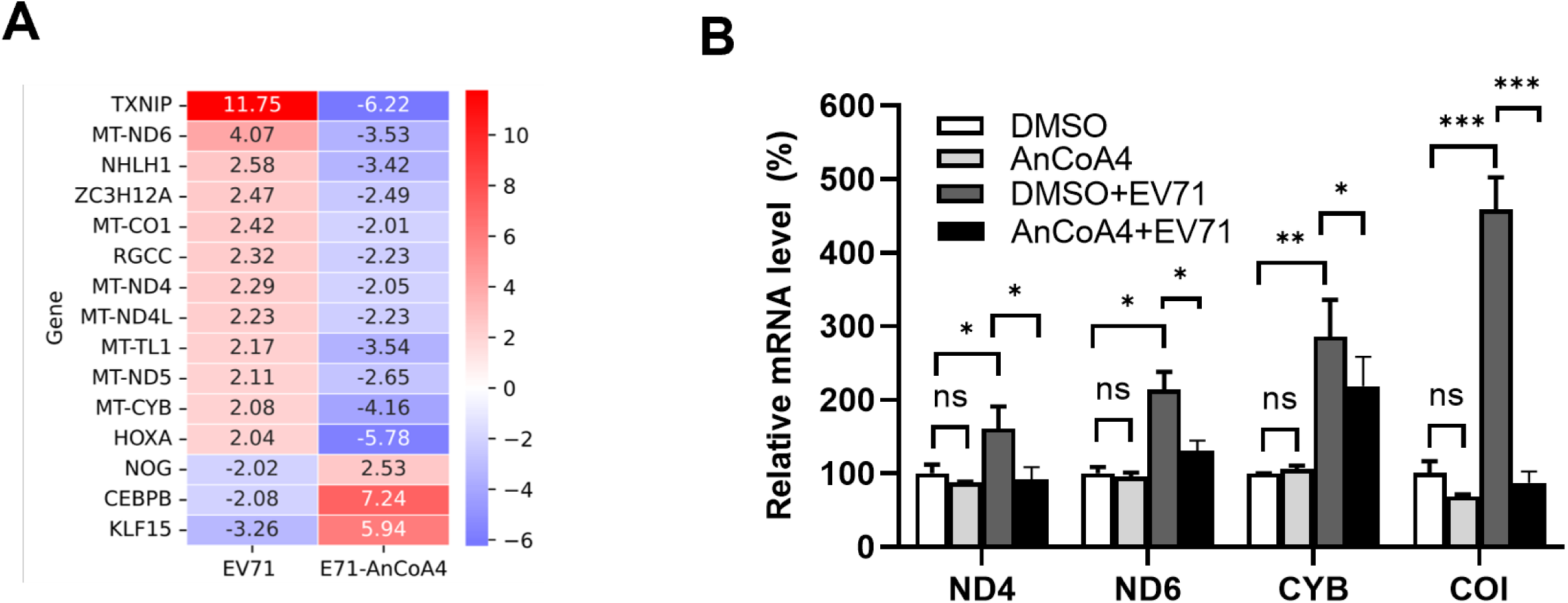
Transcriptome profiling identified mitochondrial ETC genes differentially expressed during EV71 infection.(A) Heatmap of differentially expressed genes between the DMSO- and AnCoA4-treated groups, both in the context of EV71 infection at an MOI of 0.5 for 6 h in RD cells. The numbers for each gene indicated the change folds. (B) RD cells were infected with EV71 at an MOI of 0.5 for 6 h or mock infected. The mock-infected cells or EV-infected were treated with 20 µM AnCoA4 or DMSO. Relative RNA levels of representative components of ETC complexes, as indicated, were determined through RT-qPCR by using total RNA prepared under each condition and shown as levels relative to those of mock-infected controls. RNA level was normalized with internal control GAPDH. \*\*\**p* < 0.001,\*\**p* < 0.01, \**p* < 0.05 versus the DMSO control or EV71-infected group.

**TABLE 1.** Enrichment analysis of biological processes using the DAVID in EV71-infected cells treated with AnCOA4 or left-untreated

| Biological process | Gene (Entrez gene ID)<sup>a</sup> | *p*-value<sup>b</sup> |
| --- | --- | --- |
| Electron transport coupled proton transport | MT-ND4, MT-ND5, MT-ND6, MT-CYB,<br>MT-COI | $3.15 \times 10^{-3}$ |
| Mitochondrial electron transport, NADH to ubiquinone | MT-ND4L, MT-ND4, MT-ND5, MT-ND6 | $6.68 \times 10^{-3}$ |
| ATP synthesis coupled electron transport | MT-ND4L, MT-ND4, MT-ND5, MT-ND6,<br>MT-CYB, MT-COI | $6.68 \times 10^{-3}$ |
| Response to hydrogen peroxide | MT-ND4, MT-ND5, TXNIP | $1.48 \times 10^{-1}$ |
| Mitochondrial ATP synthesis coupled proton transport | MT-ND4L, MT-ND4, MT-ND5, MT-ND6,<br>MT-CYB, MT-COI | $1.41 \times 10^{-2}$ |
| Positive regulation of transcription from<br>RNA polymerase II promoter | CEBPB, KLF15, HOXA, NHLH1, NOG,<br>RGCC, ZC3H12A | $6.69 \times 10^{-1}$ |

<sup>a</sup> Gene ID obtained from Entrez Gene resource (https://www.ncbi.nlm.nih.gov/gene/) of NCBI
<sup>b</sup> *p*-values are adjusted by Bonferroni correction method, which multiplies the raw *p*-values.

### Disruption of SOCE reduced mitochondrial ETC RNA levels

Viral 2B protein from several members of EVs causes depleted Ca^2+^ stores in the ER and Golgi apparatus, resulting in an increase in cytosolic Ca^2+^[10]. Thus, we investigated the effect of EV71 2B expression alone in HeLa cells. We identified an increase in cytosolic Ca^2+^ levels, which was significantly reduced by the treatment with either BAPTA, EGTA or AnCoA4 (Fig. 6A). Moreover, the expression of EV71 2B alone significantly upregulated the levels of ND6, CYB and COI transcripts, which were reduced by AnCoA4, as did by EV71 infection in the absence or presence of SOCE inhibitor treatment (Fig. 6B). Moreover, knockdown of either STIM1 or Orai1 resulted in marked reductions in the mitochondrial ETC RNA levels compared with those of the scramble control in the context of EV71 infection at 6 h p.i. (Fig. 6C, D); this is in agreement with those treated with AnCoA4 (Fig. 5B).

**FIG 6.**
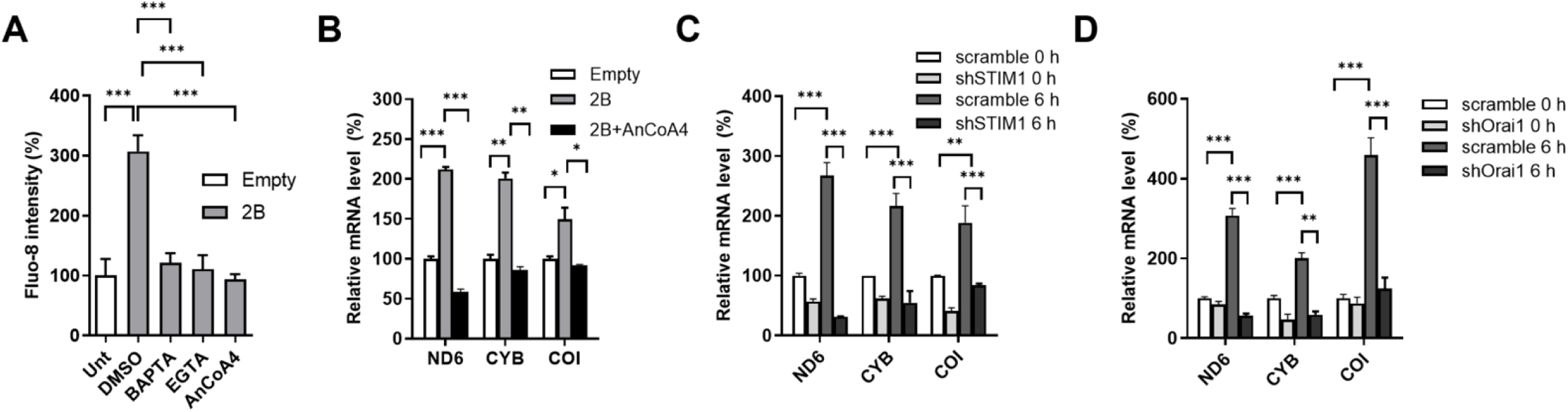
2B expression and EV71 infection induced transcriptional upregulation of mitochondrial ETC genes through SOCE machinery. (A) HeLa cells were transfected with pFLAG-CMV-2 (empty) plasmid, pCMV-FLAG-2B (2B) plasmid, or left untransfected (Unt) for 40 h. For untransfected and empty controls, media were replaced with DMSO-containing media, and incubated for 8 h. For the pCMV-FLAG-2B -transfected cells, media were replaced with DMSO-, 10 µM BAPTA-, 1.8 mM EGTA- ,or 20 µM AnCoA4-containing medium, as indicated, and also incubated for 8 h. Cells were stained with the cytosolic Ca^2+^ fluorescent dye Fluo-8. (B) HeLa cells were transfected with the empty or 2B-expressing plasmids for 48 h. HeLa cells with knockdown of STIM1 (C) or Orai1 (D), each with Scramble control, were infected with EV71 at an MOI of 0.5 for 6h. Relative RNA levels of representative components of ETC complexes, as indicated, were determined through RT-qPCR. The expression levels were related to those of mock-infected scramble controls. \*\*\**p* < 0.001,\*\**p* < 0.01, \**p* < 0.05 versus the mock-infected or DMSO control.

### EV71 infection enhanced SOCE-dependent mitochondrial respiration and ATP production

Given that the upregulation of transcripts in mitochondrial ETC complexes was observed during EV71 infection, we assessed several functional ETC parameters (indicated in the Fig. 7A) by using a live-cell Seahorse assay [35, 36] in mock-infected cells, infected cells, and infected cells treated with AnCoA4 (Fig. 7B, C). Levels of oxygen consumption rate (OCR) in EV71-infected cells were increased compared to mock-infected cells and reduced following AnCoA4 treatment (Fig. 7B). Concurrently, the relative levels of basal and maximal respiration were also increased in EV71-infected cells and reduced by AnCoA4 treatment (Fig. 7C). A similar trend of ATP production as those of the OCR and basal/maximal respiration was observed (Fig. 7C). Proton leak, spare respiratory capacity, and non-mitochondrial oxygen consumption remained unchanged. ETC begins with CI that catalyzes electron transfer from NADH to ubiquinone, which was discovered to be particularly enriched in the analysis (Table 1). We additionally investigated mitochondrial CI activity upon EV71 infection with or without AnCoA4 treatment and observed a marked elevation or reduction in CI activity, respectively (Fig. 7D). Given that ATP is the end product of the ETC, we further determined that the cellular ATP level was significantly increased following EV71 infection and reduced after AnCoA4 treatment by a luminescence assay (Fig. 7E), consistent with the result analyzed by the Seahorse assay (Fig. 7C). Moreover, viral RNA and protein levels decreased after treatment with oligomycin, an inhibitor of mitochondrial ATP synthase (complex V) [35, 36], in a dose-dependent manner (Fig. 7F, G), indicating that ATP production is essential for efficient EV71 replication. Together, these results indicate that the enhancement of the mitochondrial respiration was associated with increased mitochondrial CI activity and bioenergy production following EV71 infection at the mid-stage (6 h p.i.), with these effects likely mediated through SOCE activation.

**FIG 7.**
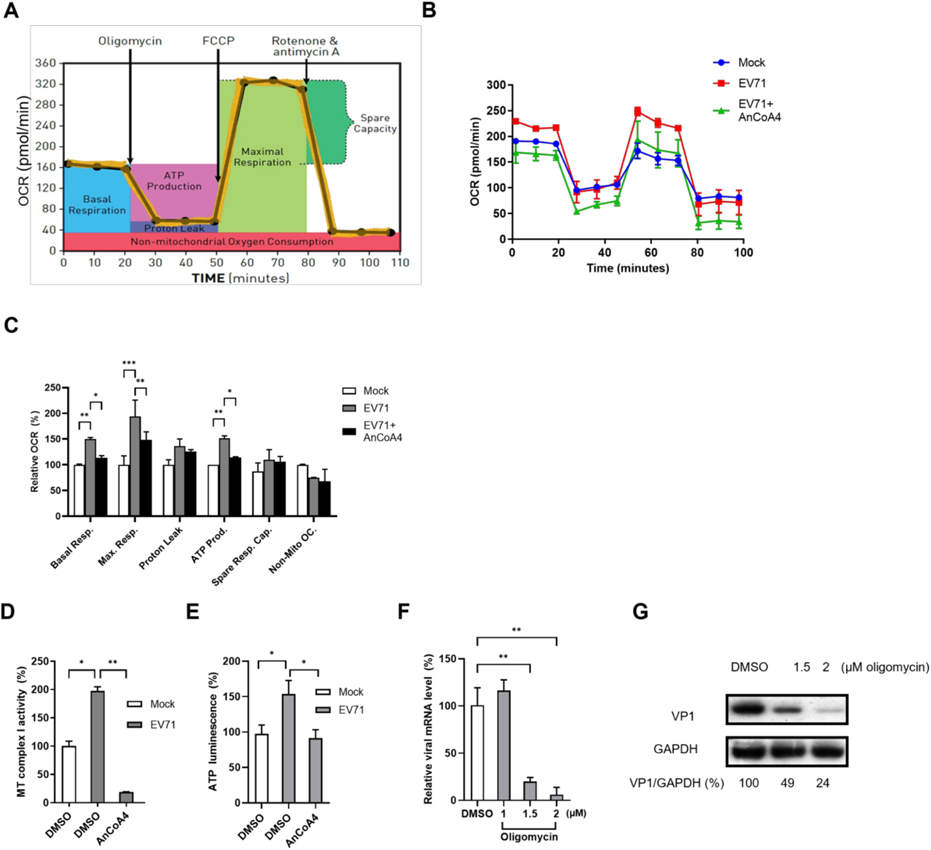
EV71-infected cells displayed increased mitochondrial respiration and ATP production through the SOCE machinery. (A) Diagram illustrating the oxygen consumption rate (OCR) measured for the parameters including basal respiration, maximal respiration, proton leak, ATP production, spare respiratory capacity, and non-mitochondrial oxygen consumption of ETC function, by using the XFe24 Seahorse analyzer; this diagram was reproduced with permission. (B) HeLa cells were mock-infected or infected with EV71 (MOI = 5) for 6 h in the absence or presence of AnCoA4 at 20 µM. Oligomycin, FCCP, and antimycin A + rotenone were injected at the times indicated in (A). (C) The six parameters were analyzed following the manufacturer’s instructions and are presented as percentages relative to the mock-infected cells. (D) Mitochondria were isolated from HeLa cells for the complex I activity assay and shown as levels relative to those of mock-infected controls. (E) Cellular ATP activity was measured by a luminescence assay and reported as levels relative to those of mock-infected controls. (F, G) HeLa cells were infected with EV71 at an MOI of 0.5 with oligomycin treated at the indicated concentrations or DMSO control. Total RNA and cell lysates were collected at 6 h p.i. for RT-qPCR (F) and immunoblots (G) , respectively. Relative RNA level was normalized with internal control GAPDH. The percentage shown below each lane represents the intensity of viral VP1 relative to that of GAPDH. Data are presented as means ± SDs. \*\*\**p* < 0.001,\*\**p* < 0.01, \**p* < 0.05 versus the mock-infected or EV71-infected groups.

### EV71 induced mitochondrial Ca^2+^ influx through SOCE

EV infections cause a progressive increase in mitochondrial Ca^2+^ during infection. In isolated mitochondrial systems, the addition of Ca^2+^ enhances the activities of complexes I, III, and IV as well as ATP production by more than two-fold [37, 38]. Therefore, we hypothesized that the upregulated levels of the complex components (Table 1 and Fig. 5A) are associated with Ca^2+^ influx into mitochondria. EV71-infected cells were stained with Rhod-2 AM (referred to as Rhod-2 hereafter) fluorescent dye to analyze the mitochondrial Ca^2+^ levels at 6 and 12 h p.i. Depletion of ER Ca^2+^ stores by TG treatment resulted in an increased level of mitochondrial Ca^2+^ compared with that in the mock control (Fig. 8A, S7). The infected cells displayed progressive elevations in mitochondrial Ca^2+^ during infection that were significantly decreased after treatment with ruthenium red (RR), a specific inhibitor of the mitochondrial Ca^2+^ uniporter (MCU) [11, 20] that allows the passage of Ca^2+^ from the cytosol into mitochondria (Fig. 8A). A similar reduction in mitochondrial Ca^2+^ levels was observed following AnCoA4 addition, implying a correlation of mitochondrial Ca^2+^ influx with SOCE activation. This notion was supported by the finding that the expression of viral 2B alone increased mitochondrial Ca^2+^ levels, which were significantly decreased following treatment with RR or AnCoA4 (Fig. 8B). Moreover, RR treatment reduced the ATP level promoted by EV71 infection (Fig. 8C), supporting the notion that elevated mitochondrial Ca^2+^ levels are associated with ATP production at 6 h p.i. Next, we analyzed a panel of representative EV serotypes, namely CVA16, CVB3, and Echo9, for the SOCE–mitochondrial Ca^2+^ axis at 6 h p.i. In all of these serotypes, we observed that either RR- or AnCoA4 reversed mitochondrial Ca^2+^ influx (Fig. 8D, F, S7), and that either treatment greatly reduced viral titers with RR treatment exhibiting a dose-dependent manner (Fig. 8E, G). These findings indicate that EV-induced SOCE activation facilitates viral replication through mitochondrial Ca^2+^ influx and consequent ATP elevation at the mid-stage of infection (6 h p.i.).

**FIG 8.**
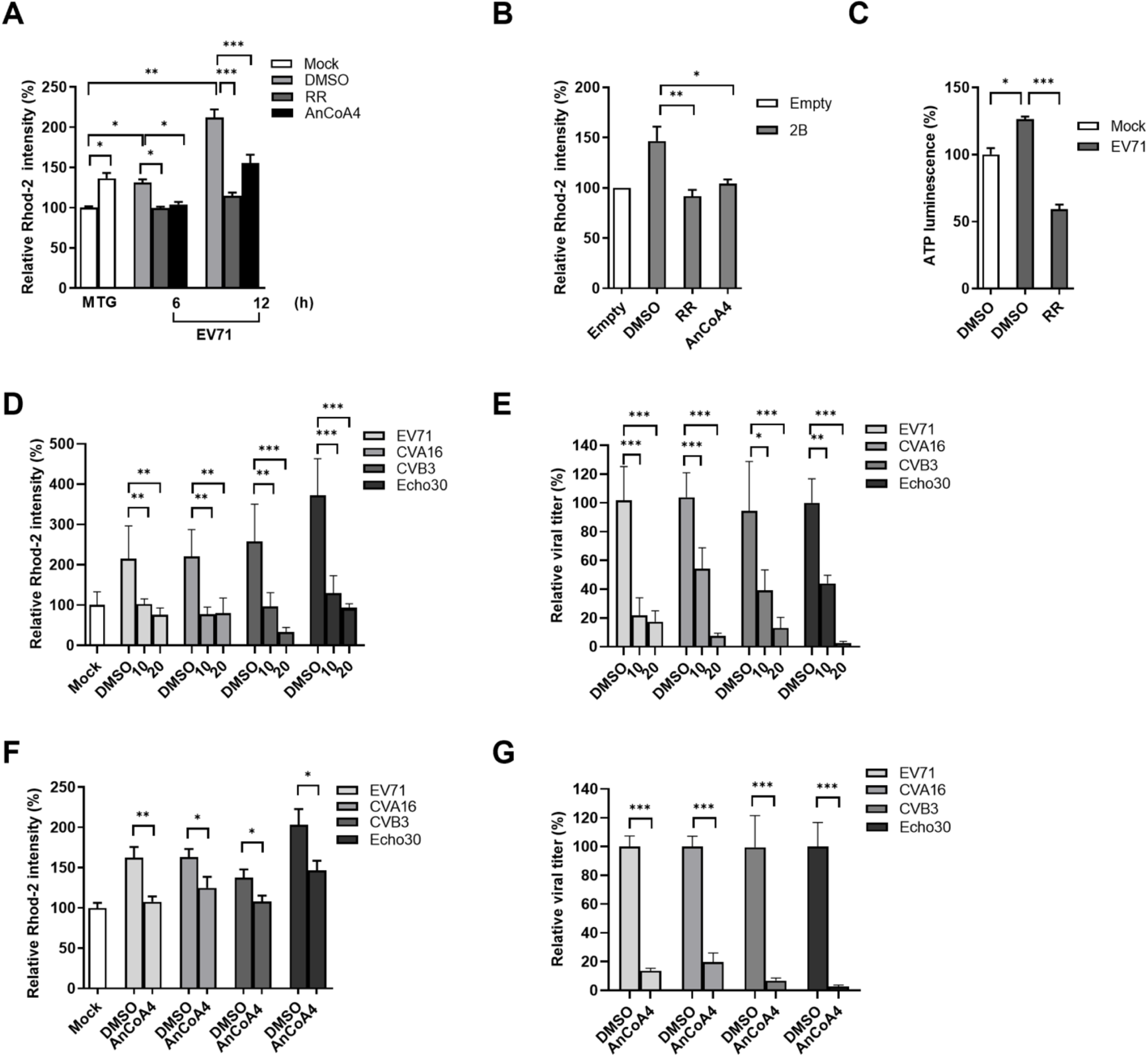
EV71 infections increased the mitochondrial Ca^2+^ influx that contributed to viral replication and ATP production at the mid-stage of the virus life cycle. (A) HeLa cells were treated with 10 µM RR, 20 µM AnCoA4, or DMSO control during EV71 infection at an MOI of 0.5 for 6 h and 12 h and were then stained with Rhod-2. TG served as a positive control for Rhod-2 stain. The relative Rhod-2 fluorescence intensity of each condition was quantified relative to that of a mock-infected control (M), and normalized by the number of cells stained with DAPI. (B) HeLa cells were transfected with an empty or 2B-expressing plasmid for 40 h. The media were replaced with DMSO, 10 µM RR, or 20 µM AnCoA4-containing media for 8 h and then stained with Rhod-2. (C) HeLa cells were infected with EV71 at an MOI of 5 for 6 h or mock-infected and treated with 10 µM RR or DMSO. Cellular ATP activity was measured and reported as levels relative to those of the mock-infected control. HeLa cells were infected with EV71, CVA16, CVB3, or Echo 30, each at an MOI of 1 for 6 h and were treated with 20 µM AnCoA4 (D, E), 10/20 µM RR (F, G), each with DMSO control, during the infection period. (D, F) Cells were stained for Rhod-2 or (E, G) total virus titer was determined by TCID_50_ assay and shown as the percentage of their DMSO controls. The experiments were conducted in triplicate, and their means ± SDs are presented. \*\*\**p* < 0.001,\*\**p* < 0.01, \**p* < 0.05 versus the mock-infected or DMSO control.

### SOCE activation is correlated with cell apoptosis and virus release during the late phase of EV71 infection

EV infections induce mitochondrial Ca^2+^ overload during the late stage of the virus life cycle, leading to cell death [8, 11, 20]. We investigated whether this Ca^2+^ influx results in mitochondrial Ca^2+^ overload and its impact on cell fate and virus production during the late phase of infection at 12 h p.i. EV71 infection markedly reduced cell viability. This reduction in cell viability was reversed by AnCoA4 in a dose-dependent manner (Fig. 9A). Consistent with this finding, the cell viability progressively decreased as the MOIs of the virus infections increased, and all of these reductions were markedly rescued by AnCoA4 treatment (Fig. 9B). Because mitochondrial Ca^2+^ overload triggers apoptotic cell death, we measured the caspase activities in the EV71-infected cells at 12 h p.i. after AnCoA4 treatment. AnCoA4 markedly reversed the EV71-induced activities of caspase -3 and -9 but not those of -8, indicating that the virus-induced activation of the intrinsic apoptosis pathway is at least partly mediated by SOCE-mediated mitochondrial Ca^2+^ overload. We next investigated whether SOCE activation and mitochondrial Ca^2+^ influx affect virus production because the release of EV from cell types such as RD cells is primarily mediated by apoptosis through the lytic mode [39]. AnCoA4 or RR was added to cultures during EV71 infection at 0.5 MOI, and the cells and their supernatants were harvested at 12 h p.i. to enable measurement of the titers of intracellular (InV) and extracellular virus (ExV), respectively. The ExV titer/total virus titer (E/T) ratio was determined to investigate the virus release efficiency. In all cases, treatment with either AnCoA4 or RR reduced the titers of ExV in a dose-dependent fashion while those of InV were not reduced (Fig 9D, F). Importantly, either compound treatment resulted in dose-dependent reduction in the E/T ratio (Fig. 9E, G), suggesting that blockade of the SOCE or mitochondrial Ca^2+^ influx led to inhibitory effects on the viral release. Virus infection at 5 MOI upon AnCoA4 treatment showed a similar trend except that the E/T ratio was higher than that with 0.5 MOI infection after treatment with 20 µM AnCoA4 (Fig. 9H, I). Together, these findings indicate that SOCE activation and subsequent mitochondrial Ca^2+^ overload play a pivotal role in the induction of apoptotic processes and facilitate the egress of virus progeny at the late stage (12 h p.i.) of an infection cycle.

**FIG 9.**
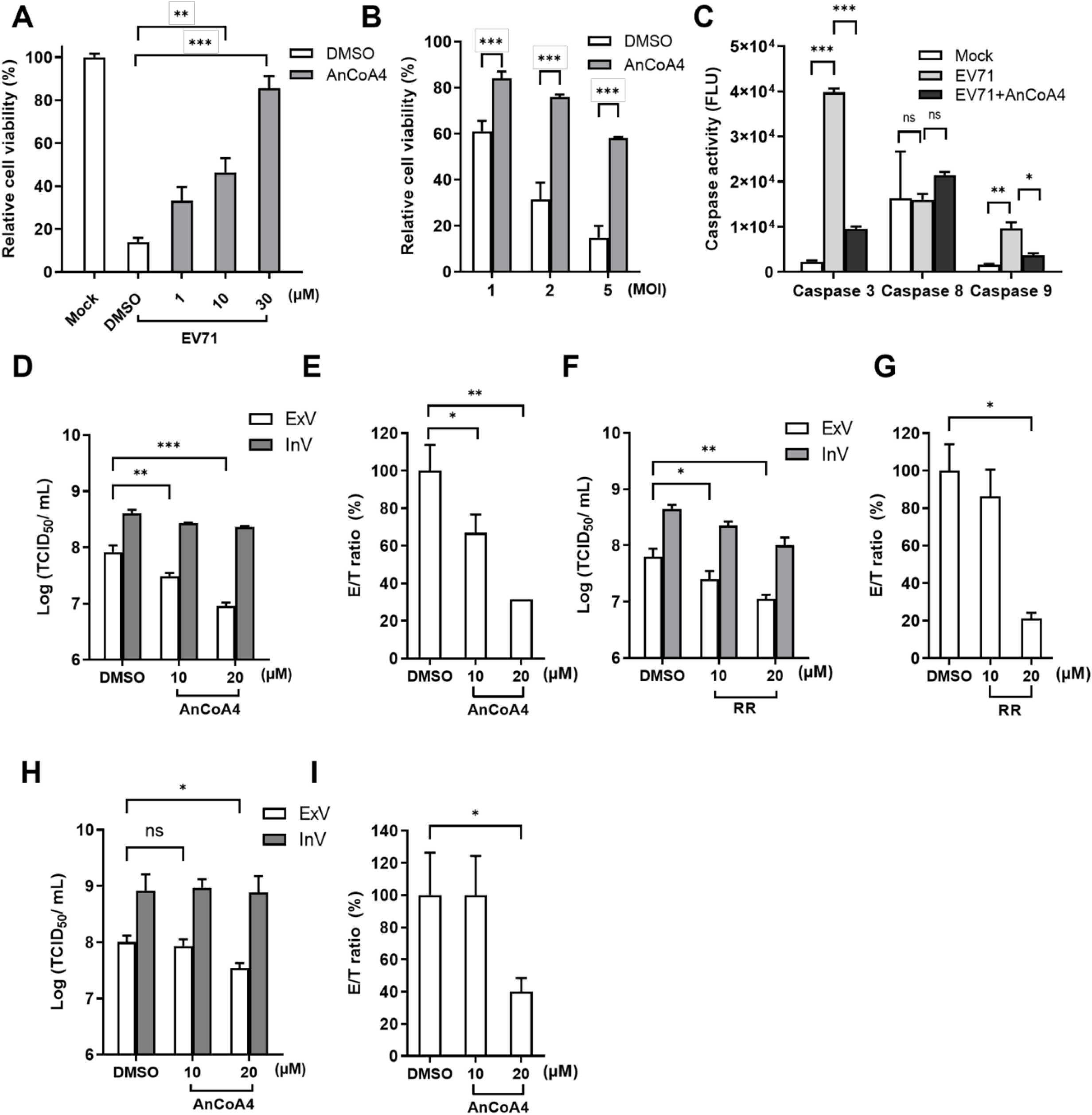
The SOCE inhibitor AnCoA4 attenuated EV71-induced apoptosis and the viral release at the late stage of infection. (A) RD cells were mock-infected or infected with EV71 at an MOI of 5 for 12 h. The infected cells were treated with the indicated concentrations of AnCoA4 or DMSO control. (B) EV71 was used to infect RD cells at MOIs of 1, 2, and 5 for 12 h, and cells were treated with 10 µM AnCoA4 or DMSO control. Cell viability was measured using the MTS assay and compared with that of mock-infected controls . (C) RD cells were mock-infected or infected with EV71 at an MOI of 1 for 12 h and treated with 20 µM AnCoA4 during infection or DMSO control. Caspase-3, -8, and -9 activities were measured using the caspase fluorometric assays. RD cells were infected with EV71 at an MOI of 0.5 for 12 h and treated with AnCoA4 (D) or RR (F), both at 10- and 20-µM, during the infection period. At 12 h p.i., the supernatant was collected for the extracellular virus (ExV), and cells were trypsinized for preparing the intracellular virus (InV). The titers of ExVs and InVs in each group were determined using the TCID_50_ assay (D, F). For each condition, the relative ratio of the ExV titer over the total virus titer, compared with that of untreated control, is represented by the E/T ratio (%). (H) RD cells were infected with EV71 at an MOI of 5 for 12 h and treated with AnCoA4 at 10- and 20-µM. At 12 h p.i., the supernatant was collected for the extracellular virus (ExV), and intracellular virus (InV). The titers of ExVs and InVs in each group were determined using the TCID_50_ assay (D, F, H). For each condition, the relative ratio of the ExV titer over the total virus titer, compared with that of untreated control, is represented by the E/T ratio (%) (E, G, I) and indicates the virus release efficiency. Data represented the means of triplicated experiments and the SD Data represented the means of triplicated experiments and the SD. \*\*\**p* < 0.001,\*\**p* < 0.01, \**p* < 0.05 versus the DMSO control.

### Mouse intestinal organoids infected with EV71

Given that all studies above were conducted using permissive cell lines, that is, RD and HeLa cells, we investigated Ca^2+^ signal–dependent viral replication in a more physiologically relevant cell culture system. We established a mouse intestinal organoid culture derived from isolated intestinal stem cell–containing crypt fractions to evaluate intestinal pathophysiology [22, 28, 40]. Single intestinal stem cells can be cultured into three-dimensional enteroids comprising all major differentiated cell types that can be found in the native intestinal organoid (Fig. 10A). However, EV infections are restricted by species barriers; human cells are susceptible to such infections, whereas mouse cells are typically resistant to them [41]. On the other hand, a mouse-adapted EV71 MP4 strain was demonstrated to efficiently infect mouse cell cultures and a mouse model [26, 42].

**FIG 10.**
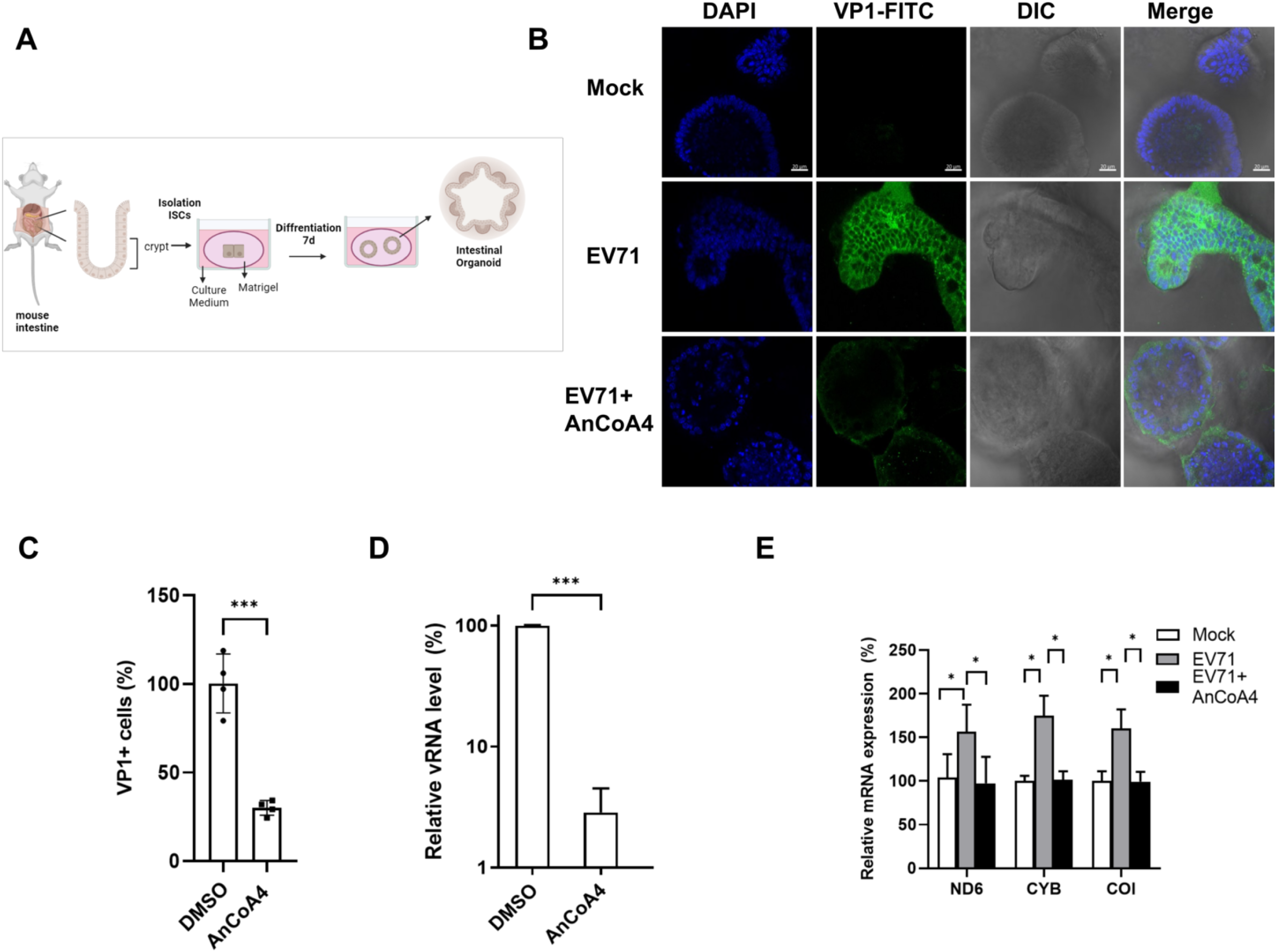
EV71 replication and virus-upregulated ETC RNA were reduced by AnCoA4 in the mouse intestinal organoid model. (A) Schematic representation of the three-dimensional mouse intestinal organoid culture. (B) Mouse intestinal organoid cultures seeded in Xfe 24-well dishes were inoculated with 200µl EV71 MP4 strain (4 *×* 10^8^ TCID_50_/mL) for 48 h and treated with 20µM AnCoA4 during the infection period or DMSO control. Immunofluorescence assay was conducted, with viral VP1 antigen detected using the mouse anti-EV71 Ab (green) and the nucleus stained with DAPI (blue). (C) The relative VP1 positive (+) antigen expression of MP4-infected, DMSO-treated, and MP4 infected, AnCoA4-treated cells were quantified relative to the number of cells stained with DAPI. Total RNA was prepared, and viral RNA (D) and the indicated mitochondrial ETC RNA (E) were measured through RT-qPCR and presented as levels relative to those of the mock-infected controls (%). Data are presented as means ± SDs. \*\*\**p* < 0.001,\*\**p* < 0.01, \**p* < 0.05 versus the DMSO control.

Thus, we used a mouse-adapted EV71 MP4 strain to inoculate our mouse intestinal enteroid cultures in the absence or presence of AnCoA4. Immunofluorescence staining was conducted, and the levels of viral antigen were analyzed through confocal microscopy. The viral antigen signals were well discernible in the infected-cells and significantly reduced after AnCoA4 treatment (Fig. 10B, C). In addition, the relative levels of viral RNA and representative ETC ND6, CYB and COI RNAs were analyzed. A similar trend of alteration in the viral RNA was observed as those of the viral antigen (Fig. 10C, D). The mouse ETC RNA levels were substantially increased following EV71 infection relative to that of the mock-infected control; the levels significantly declined upon treatment with AnCoA4 (Fig. 10E).

## DISCUSSION

In the present study, we aimed to investigate the molecular mechanism underlying the Ca^2+^ flux and its impact on the cell fate and EV life cycle. We showed that the SOCE machinery likely mediated the increase in cytosolic Ca^2+^ and viral replication during the mid-phase of EV71 life cycle (Fig. 1–4). Moreover, we demonstrated that the ETC complex genes were enriched through SOCE during the mid-stage of infection (Table 1, Fig 5A).

This resulted in alterations in the mitochondrial ETC CI activity, ATP production and cellular respiration (Fig. 5, 6, 7) through SOCE and mitochondrial Ca^2+^ influx, and contributed to viral replication (Fig. 6, 8). The SOCE–mitochondrial Ca^2+^ influx axis also plays a role, at least in part, in triggering apoptosis and subsequent viral release due to mitochondrial Ca^2+^ overload during the late stage of virus infection (Fig. 9).

We started with investigating the potential contribution of the enhanced cytosolic Ca^2+^ to different stages of the viral life cycle. It was demonstrated that regardless of the infection doses, EV71 replication upon the treatments of BAPTA, EGTA or AnCoA4 showed the delay kinetics relative to that of untreated control, with the delay most pronounced at 4 h and 6 h p.i. (Fig. 1B, 1D, 2C, 2E, S3). This is consistent with the finding that AnCoA4 treatment between 3-6 h after viral adsorption, but not other periods assessed, resulted in a reduced viral titer (Fig. 2F, G). It is speculated that viral-enhanced mitochondrial function and ATP production facilitate virus replication during the mid-stage of replication. Once the infection passes the virus replication stage highly required for ATP and proceeds to 9 h and 12 h p.i., the differences in viral titers between the AnCoA4-treated and DMSO control groups significantly decreased (Fig. 2C, E).

Nevertheless, virus release was significantly impaired regardless of the inoculated virus doses at 12 h p.i., likely due to inhibition of the mitochondrial Ca^2+^ overload and the consequent proapoptotic state by AnCoA4 (Fig. 9H, I). Indeed, these data are in agreement with the report that recombinant coxsackievirus B3 carrying the mutant 2B also exhibited a delay in viral replication at the earlier stage and a defect in virus release and that virus release was greatly hampered when the infected cells were cultured in the Ca^2+^-free medium compared with that in the Ca^2+^-containing medium [10]. This proposed mechanism (Fig. 11) is supported by our finding that the expression of EV71 2B alone resulted in SOCE-dependent increases in the ETC transcripts and mitochondrial Ca^2+^ levels (Fig 6B, 8B).

**FIG 11.**
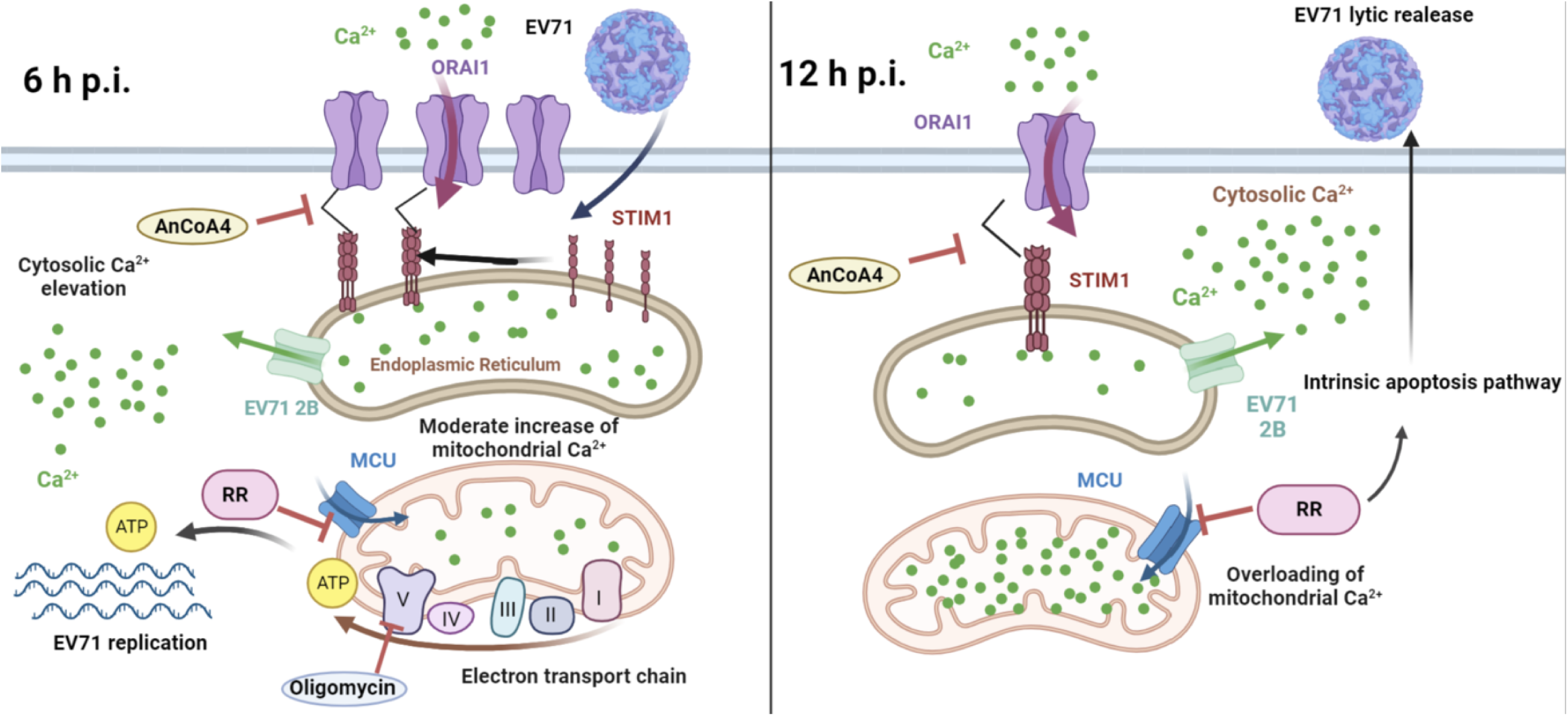
Proposed mechanism through which EV71-induced cytosolic Ca^2+^ elevation affects the virus life cycle through SOCE activation and mitochondrial Ca^2+^ influx. EV71 2B caused ER Ca^2+^ depletion, resulting in SOCE activation. At 6 h p.i, cytosolic Ca^2+^ levels increased, causing a moderate elevation in mitochondrial Ca^2+^ and subsequent increases in mitochondrial electron transport chain complex (I through IV) activity and ATP generation for viral replication. At 12 h p.i, overloading of mitochondrial Ca^2+^ promoted viral release through apoptotic cell death triggered by the intrinsic apoptosis pathway. AnCoA4 and RR reduced the levels of cytosolic Ca^2+^ and mitochondrial Ca^2+^, reducing viral replication and virus release, respectively. Oligomycin targets the mitochondrial complex V (ATP synthase), leading to inhibition of viral replication. RR = ruthenium red. MCU = mitochondrial Ca^2+^ uniporter

Individual expression of EV 2B is known to induce the influx of extracellular Ca^2+^; however, the precise mechanism has not been defined [7, 10]. In this study, we clearly showed that knockdown of either STIM1 or Orai1 resulted in reduced levels of the virus-induced cytosolic Ca^2+^ and mitochondrial ETC RNA as well as those of viral protein and viral titers (Fig. 3A-C, Fig. 6C, D). These data indicate that the STIM1-Orai1 machinery contributes to the extracellular Ca^2+^ influx into the cytosol during EV infection, consistent with the effects caused by AnCoA4 treatments (Fig. 2, 5). The STIM1–Orai1 machinery has been reported to mediate the replication of some viruses. The activation of the STIM1 and Orai1 channels promotes virion assembly and budding during Ebola or Marburg virus infection [43]. The replication capacity of certain viruses, such as dengue virus and rotavirus [44, 45], relies on Ca^2+^ influx through the STIM1–Orai1 system. The hepatitis B virus HBx protein directly binds to and modifies STIM1–Orai1 complexes, regulating Ca^2+^ levels and thereby promoting viral replication by modulating mitochondrial Ca^2+^ uptake [46]. However, the downstream signaling pathways responsible for their replication remain unknown. Some viruses inactivate SOCE to reduce immune responses and facilitate viral replication. For example, a study reported that the downregulation of type I interferon through Orai1 depletion facilitated SARS-CoV-2 infection [47]. In the current study, we determined that SOCE participates in two steps of the virus life cycle, namely viral replication and virion release, through progressive mitochondrial Ca^2+^ influx. This represents a novel mechanism involving SOCE-mediated virus replication.

The findings of our RNA-seq/bioinformatic analysis revealed that a panel of components of mitochondrial ETC complexes were enriched. Because the primary function of mitochondria is to generate ATP through oxidative phosphorylation, we investigated the function correlations between these components and determined that EV71 infection increased mitochondrial ETC CI activity, ATP levels, and cellular respiration. These effects were significantly attenuated following the blockade of SOCE or mitochondrial Ca^2+^ influx by AnCoA4 or RR, respectively. These findings are consistent with those of previous studies indicating that a moderate increase in mitochondrial Ca^2+^ promotes the activities of the ETC CI, CIII, and CIV as well as ATP production [37, 38, 48]. Regarding the role of increased ATP in EV replication, studies have demonstrated that EV 2C plays a crucial role in virus replication by acting as an ATP-dependent RNA helicase for virus RNA replication. Moreover, EV71 infection induces mitochondrial clustering and recruitment, suggesting localized ATP production occurs at the virus replication site. Prior to this study, some viruses, including cytomegalovirus, dengue virus, and vaccinia virus, have been reported to promote mitochondrial ETC function and subsequent energy output to support virus replication [35, 36, 49]. Although the mechanisms underlying this phenomenon remain largely unknown for dengue virus and vaccinia virus, the cytomegalovirus protein pUL13 has been demonstrated to interact with mitochondrial cristae to enhance ETC function [36]. On the other hand, some other viruses undergo their replication without upregulating ETC activity. This is best known for hepatitis C virus and influenza viruses whose ATP-dependent replication mainly rely on the glycolysis pathways rather than ETC activity [50, 51, 52]. This is in marked contrast to our RNA-seq analysis where the mitochondrial ETC transcripts, but not those in the glycolysis pathways, were enriched (Table 1, Fig S6). To the best of our knowledge, this is the first study to report that EV71 infection augments ETC activity and ATP production to support virus replication and that it employs distinct mechanisms involving SOCE activation and mitochondrial Ca^2+^ influx.

Studies on a diverse group of viruses, including EVs, have revealed that viruses induce substantial and persistent increases in mitochondrial Ca^2+^ levels, leading to apoptotic cell death, especially during the late stages of the infection cycle. In addition to being localized in the ER, the EV71 2B protein was demonstrated to localize to mitochondria, thereby directly binding to and activating the proapoptotic protein Bax [53].

This interaction ensures the induction of cell apoptosis, potentially exacerbating the intrinsic apoptosis driven by mitochondrial Ca^2+^ overload, as demonstrated in this study. Other mechanisms, including EV71-induced, caspase-independent activation of the apoptosis-inducing factor, may contribute to apoptosis, maximizing cell death and virus release.

Human intestinal organoid models have been established and used to study EVs, including EV71 [54, 55, 56]. In this study, we used a mouse intestinal organoid culture system developed from an inbred mouse origin (Fig. 10A). This system offers a stable, homogenous and continuous supply of organoids [22, 28, 40], whereas a human model does not [57]. Compared with infections by human EV71 strain without mouse-adaptation, the use of a mouse-adapted MP4 strain greatly enhanced the infection efficiency in mice [26, 42] and in the mouse intestinal organoid culture (our unpublished data). We used the mouse-adapted EV71 strain MP4 for infection, and viral antigen was well discernible in the mouse intestinal organoids in a SOCE-dependent manner (Fig. 10B, C). This is consistent with the report that SOCE activation could be demonstrated in a human intestinal organoid culture [58]. In addition, we observed a concurrent increase in the levels of mouse ETC ND6, CYB and COI transcripts along with EV71 RNA, all in a SOCE-dependent manner (Fig. 10D, E). These findings indicate that our mouse intestinal organoid system provides a valuable and physiologically relevant platform for in-depth study of virus-induced alterations in host Ca2+ signaling.

Our study demonstrated that the SOCE- and mitochondrial Ca^2+^ influx-dependent virus replication can be recapitulated with infections involving other representative EV serotypes (Fig. 8E, G). This led us to speculate whether spatial and temporal control of Ca2+ signals, as observed in EV infections, is a common manifestation in viral replication. If this speculation is confirmed, host channels that support these Ca2+ signals, such as STIM1 and Orai1, may represent novel targets for the development of broadly effective host-directed antiviral therapeutics.

## ACKNOWLEDGMENTS

This study was supported in part by the Professor Tsuei-Chu Mong Merit Scholarship (grant 40719001), NYMU-FEMH Joint Research Program (grant 110DN40), and the Ministry of Science and Technology, Taiwan, ROC (111-2320-B-A49-026-).

